# Chronically implantable LED arrays for behavioral optogenetics in primates

**DOI:** 10.1101/2020.09.10.291583

**Authors:** Rishi Rajalingham, Michael Sorenson, Reza Azadi, Simon Bohn, James J DiCarlo, Arash Afraz

## Abstract

Challenges in behavioral optogenetics in large brains demand development of a chronically implantable platform for light delivery. We have developed Opto-Array, a chronically implantable array of LEDs for high-throughput optogenetic perturbation in non-human primates. We tested the Opto-Array in the primary visual cortex of a macaque monkey, and demonstrated that optogenetic cortical silencing by the Opto-Array results in reliable retinotopic visual deficits on a luminance discrimination task.

---

A key goal in systems neuroscience is to uncover the specific neural mechanisms that underlie behaviors of interest. To this end, perturbation tools such as pharmacological, electrical and optogenetic stimulation and inhibition of neural activity, have been critical to test the causal role of neural activity in different brain sub-regions in various behaviors. In particular, optogenetic perturbations, whereby light-sensitive ion channels/pumps ^1,2^ are embedded in the membrane of genetically targeted neurons to modulate their activity via delivery of light, offer tremendous promise for neuroscience research, affording the ability to both drive and inhibit neural activity with precise temporal delimitation and cell-type specificity.

While the toolbox of optogenetic methods has been widely and successfully used in rodent brains, this method is still relatively under-developed for non-human primates (NHP) such as rhesus macaques, an animal model with a large brain, expressing highly sophisticated sensory, motor and cognitive behaviors. Indeed, only a handful of studies in NHPs show behavioral effects of optogenetic perturbation, across sensory, motor, and cognitive domains despite tremendous interests ^3–10^. The dearth of documented behavioral impacts using optogenetics may stem from several problems, including difficulties of successful genetic targeting of neurons and of delivering sufficient light to perturb those neurons in the primate brain. A typical primate optogenetic experiment consists of first injecting a viral opsin acutely in the brain, either in a sterile surgery or through an implanted recording chamber. Following viral expression in the targeted cortical tissue, light is delivered through an optical fiber, acutely inserted into the brain coupled with a recording electrode ^11^, that is driven by an external LASER or LED light source.

There are two major problems with light delivery through an optical fiber. First, the acute nature of optical fiber experiments limits the number of experimental conditions and data trials, as the fiber cannot return to an exactly similar position across multiple days (hereafter termed “chronic-repeatability”). Second, given the size and shape of optical fibers, each penetration comes with a significant cost of tissue damage and risk of hitting small arteries on the fiber path (hereafter termed “tissue-damage”). This severely limits the number of practical fiber penetrations, and thus constrains the number of variables, experiment conditions and trial counts available to the scientist. Moreover, the damage associated with fiber penetrations constrains the maximum diameter of the fiber, thus significantly limiting the cortical surface area that can be illuminated (hereafter termed “illumination-scale” and “illumination-resolution”). This is a considerable limitation, particularly when working with large brains.

There have been several attempts to innovate on this typical optical fiber-based experimental approach. First, by sharpening the tip of the fiber, it is possible to increase the cone of illumination while maintaining a small fiber diameter ^12–14^, but this gain in illumination-scale is relatively modest and the approach remains an acute protocol, thereby not addressing problems related to chronic-repeatability and tissue-damage. Direct illumination of cortex through transparent artificial dura has been successfully used to bypass the problems of optical fibers ^15–17^. This approach is highly promising, as it allows for flexible illumination-scale and resolution, mitigates the aforementioned tissue-damage problems, and could be used in a chronic manner to solve problems related to chronic-repeatability. Moreover, this approach can be coupled with red-shifted opsins to further enhance illumination scale ^18^. However, it poses other challenges, including the risk of infection and is limited to use in brain subregions that permit direct optical access to the brain surface. Chronically implanted illumination methods could in principle address many of these problems ^19,20^, as they allow reliable targeting of the same cortical position across multiple days, and do not pose any safety issues related to tissue-damage from acute probe insertions or infection from open chambers. However, given difficulties arising from the number of independently controlled illumination sources, no existing chronic illumination device is currently capable of both large-scale and high-resolution illumination.

To address this problem and improve the utility of optogenetics in non-human primates, we have developed Opto-Array (Blackrock Microsystems), a chronically implantable array of LEDs for light delivery in optogenetic experiments in primates. This tool harnesses the advantages of existing optogenetics — the precise spatial and temporal control of genetically specific neural activity — but offers three additional key advantages. First, the chronic nature of this perturbation tool enables highly stable experimental perturbation of the same neural population over months, thus dramatically increasing the scale (both number of trials, but also number of unique conditions) and throughput of current causal experiments. Second, the 2D matrix array configuration of LEDs enables the flexible perturbation of a large cortical region at fine resolution. Illuminating individual LEDs corresponds to focused perturbation of specific mm-scale columns, whereas simultaneously illuminating (arbitrary patterns of) multiple LEDs corresponds to perturbation of larger cortical areas (currently up to 5mmx5mm for each array). Third, the Opto-Array provides a safe and easy alternative to acute methods as well as direct illumination methods for light delivery, minimizing the tissue damage that results from inserting large optical fibers into the cortical tissue, as well as the risk of infection associated with open cranial windows and chambers. Additionally, the Opto-Array includes an on-board thermal sensor to monitor heating (and potential damage) of the cortical tissue from light delivery. The shortcomings of the optical array in its current format include its limitation to surface areas of the cortex (although implantation in large sulci and areas without direct visual access is possible, e.g. over inferior temporal cortex) and its lack of neural recording probes. Given the current challenges in behavioral optogenetics in large brains, we designed the first generation of Opto-Array specifically for behavioral experiments.

As shown in Figure 1A, each LED array consists of a 5×5 printed circuit board (PCB) grid with 24 LEDs (Green 527nm LEDs were used here) and one thermal sensor for monitoring tissue heating from electrical power. Each LED is 0.5mm×0.5mm, with 1mm spacing between LEDs. The PCB and LEDs are encapsulated within a thin (<0.5mm; total array thickness of 1.5 mm) translucent silicone cover. The LED array is designed to be chronically implanted directly on the cortical surface by suturing the silicone encapsulation onto the dura mater (Figure 1F). The LED array is powered through a thin gold wire bundle terminating on a Cereport pedestal connector that is implanted on the skull surface. Together, this implant allows for the delivery of light to a large region of the cortical surface with high spatial and temporal precision and stability over months of data collection.

**Figure 1.**
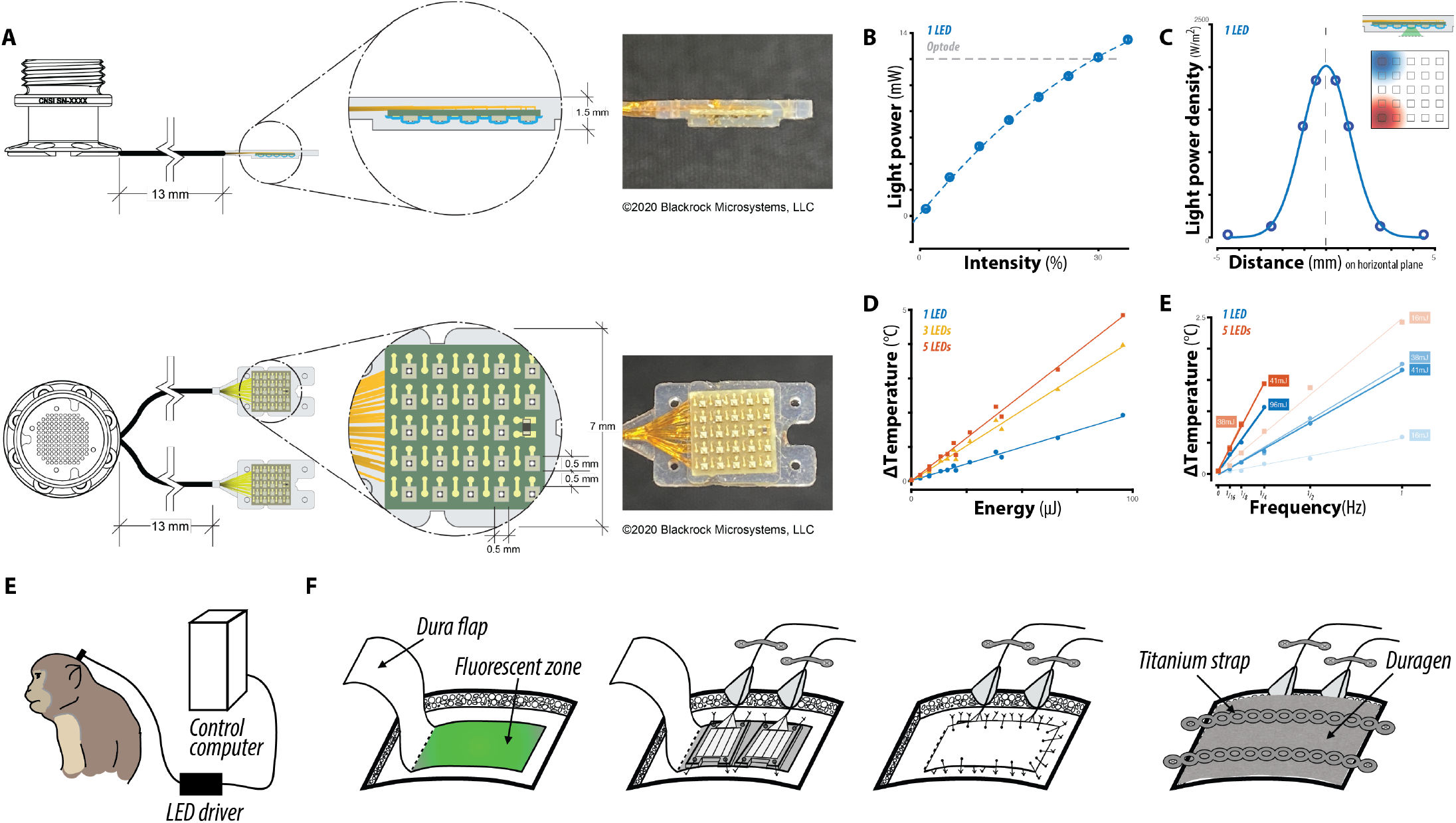
(a) Schematic of the Opto-Array design, consisting of a 5×5 grid with 24 LEDs and one thermal sensor on a PCB encapsulated in a thin translucent silicone cover. The array is designed to be chronically implanted directly on the cortical surface, by suturing the silicone encapsulation onto the dura mater (see inset). The LED array is powered through a thin gold wire bundle terminating on a Cereport pedestal connector that is implanted on the skull surface. (b) Light power output for individual LEDs as a function of the input intensity (controlled via input voltage). The horizontal dashed line corresponds to average power output of optrodes that have successfully yielded measurable behavioral effects in monkeys. (c) Spatial density of light power on the horizontal plane, at a transverse distance of <1mm from the surface of the LED. The spatial spread of light power over the horizontal plane is largely constrained to a millimeter-scale region, ensuring that activating individual LEDs yields distinct light patterns on the cortical surface (see inset). (d) Average maximum increase in temperature, measured from on-board thermal sensor, from activating different groups of LEDs as a function of the input energy (combining electrical power and illumination duration). (e) Corresponding average increase in thermal sensor response from varying the temporal frequency of activation. Measurements in (d) and (e) correspond to conservative upper bounds for the corresponding temperature change on the cortical surface, given heat transfer through the OptoArray’s silicone encapsulation. (f) Schematic of primate behavioral experiment, with chronically implanted Opto-Array sutured onto the dura mater and connected via Cereport pedestal to an external LED driver. (g) Schematic of surgical implant of Opto-Array showing suturing of Opto-Array onto dura flap, sutured closing of dura mater, and titanium strap cover on craniotomy.

We first characterized the photometric properties of the Opto-Array for direct comparison with an alternative light delivery method for optogenetic perturbation. Figure 1B shows the total light power output of a given LED, as a function of applied voltage plotted as percentage of the maximum voltage. Individual LEDs operating at 30% intensity match the power output of optical fibers that have successfully yielded measurable behavioral effects in monkeys (10-15mW). Figure 1C shows the spatial density of light power on the horizontal plane, at a transverse distance of <1mm from the surface of the LED. While light delivered from LEDs is not collimated (as for a LASER), the spatial spread of light power over the horizontal plane is sufficient to distinguish between neighbouring LEDs (half-max-full-width HMFW=2.6mm).

Next, we characterized the thermal response of the Opto-Array using the on-board thermal sensor. We note that this measurement is a conservative upper bound for the corresponding temperature change on the cortical surface, as each 1° increase measured by the thermal sensor corresponds approximately to an increase of 0.02° to 0.26° on the external surface of the Opto-Array (see Methods). We aim to limit illumination-driven tissue heating because increasing the cortical temperature above 4°C can induce tissue damage (Galvan et al. 2017). Figure 1D shows the average increase in thermal sensor response from activating different groups of LEDs as a function of the illumination energy (combining electrical power and illumination duration), at a fixed low frequency of activation. Figure 1E shows the corresponding average increase in thermal sensor response from varying the temporal frequency of activation. Together, these data demonstrate that the Opto-Array can reliably measure heating caused by LED illumination, also that typical experimental usage results in heating significantly below the risks of tissue damage.

We then tested the efficacy of the Opto-Array in-vivo in a primate behavioral experiment. As a proof of concept, we investigated the causal role of mesoscale subregions in the primary visual cortex (V1) of a macaque monkey in the context of a two-alternative-forced-choice (2AFC) luminance discrimination task (see Methods, Figure 2A). Briefly, we trained a monkey to report the location of a visual target stimulus based on its luminance, in the presence of a distractor stimulus. By varying the relative luminance of the two stimuli, we systematically varied the task difficulty. As shown in Figure 2C, the monkey’s performance varied systematically with the task difficulty as expected, with increased probability of choosing a region of the visual field with increased visual signal (the difference in luminance between the stimulus in the region and the stimulus outside the region). Stimuli were presented at randomly selected locations in the visual field within a fixed range of eccentricity, resulting in a disc of tested visual space. We then implanted two LED arrays over a dorsal region of the right V1 that was previously infected with AAV8-CAG-ArchT. Viral expression and neuromodulation were verified via a small number of acute optrode experiments (Figure S1). Given the functional organization of V1, behavioral effects from perturbing this cortical region are expected to be spatially constrained on the visual field (target ROI, contralateral lower visual field, Figure 2D). Given the spatial symmetry of the task, we additionally expect an equal and opposite behavioral effect in the radially opposite position in the visual field.

**Figure 2:**
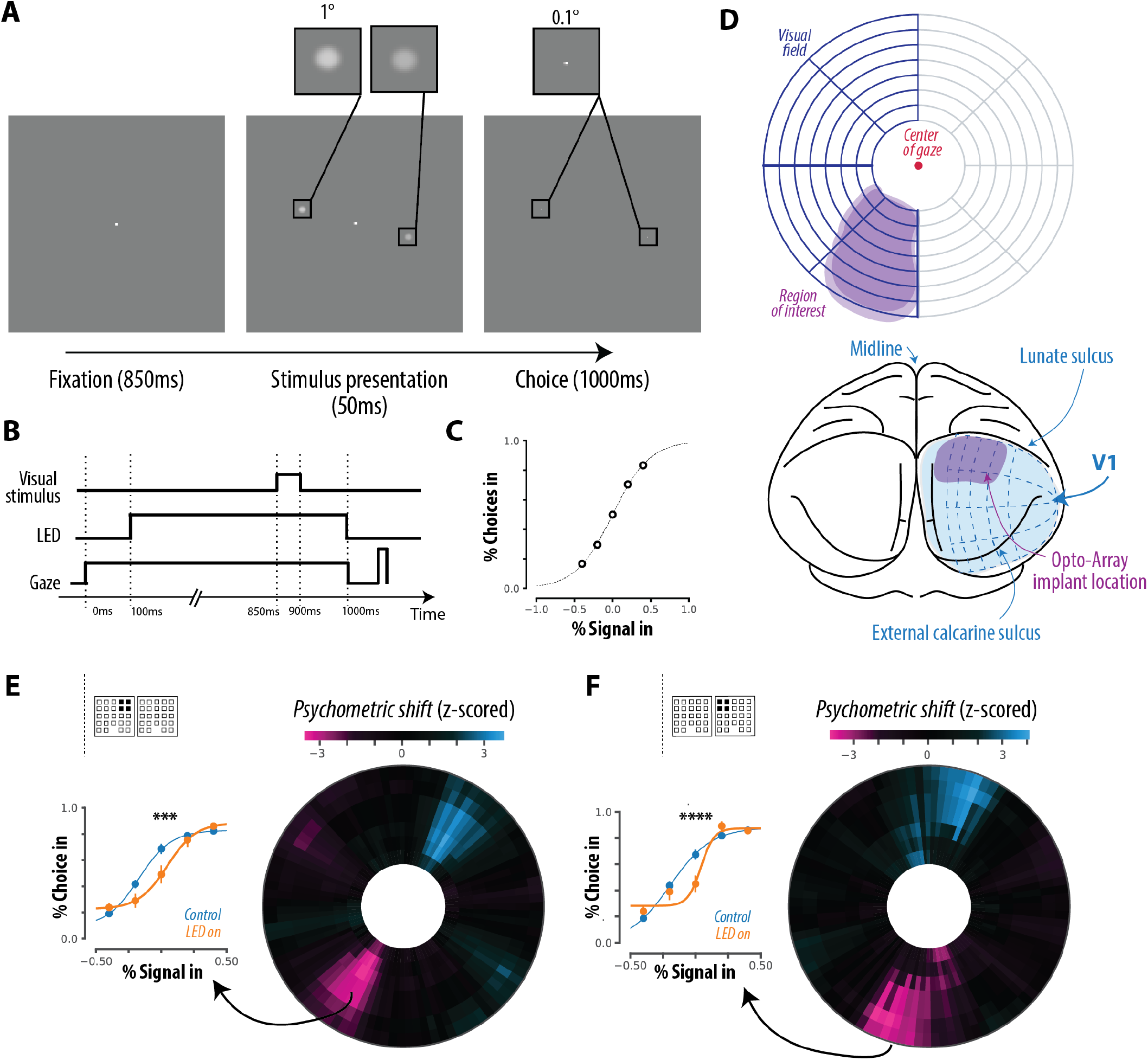
(a) Behavioral paradigm for luminance discrimination task. Each trial of the behavioral task consisted of a fixation period, during which one (or none) of the LEDs were preemptively activated on a random proportion of trials. Following fixation, two sample stimuli were briefly presented at random radially opposite locations in the visual field. The task required the subject to make a saccade to a target location defined by the brighter of the two sample stimuli. The location and relative luminance of the stimuli was randomly assigned for each trial. By varying the relative luminance of the two sample stimuli, we systematically varied the task difficulty. (b) The time course of the behavioral paradigm. The LED activation was timed to completely overlap the stimulus-related activity in V1. (c) Control behavior from the animal. (d) Correspondence between spatial organization of V1 cortex (bottom) and the visuospatial organization of the visual field (top). Behavioral effects from perturbing the Opto-Array implant region are expected to be spatially constrained to a target ROI, shown in purple. Given the spatial symmetry of the task, we additionally expect an equal and opposite behavioral effect in the radially opposite position in the visual field. (e,f) For two different example LED conditions (see insets for location of activated LEDs), z-scored psychometric shift maps are shown, with raw data and fitted psychometric curves from the target regions shown on the left.

We measured the monkey’s behavior on the luminance discrimination task, comparing illumination versus control trials. To maximize both the spatial spread and power of light, we activated groups of four neighboring LEDs simultaneously, and interleaved four such groups. Given the chronic nature of this tool, we collected behavioral data over several sessions while activating LEDs on a small portion of trials (20%). Pooling over all LED conditions and over the entire ROI, we observed a reliable behavioral effect of LED illumination even at this coarse scale, in the form of a statistically significant psychometric shift for a spatially restricted subregion of the visual field encompassed within the ROI (*p=4*.*75e-4*, Figure S1E). We then analyzed the corresponding effects over different LED conditions and different subregions within the target ROI. Figure 2C shows the psychometric shift maps for two different example activation conditions (each of four neighbouring LEDs); the insets show the locations of each of the four activated LEDs. Each map shows a reliable behavioral shift at subregions of the visual field encompassed within the ROI (*p=1*.*84e-4, 3*.*67e-5*), where each effect is spatially restricted to a distinct subregion of the visual field encompassed within the ROI. These results demonstrate that, even in spite of the weak viral expression and photo-suppression of neural activity we observed here, illumination from the Opto-Array results in reliable spatially-restricted behavioral effects, validating this tool for behavioral experiments with optogenetic perturbation. The specific spatial illumination parameters necessary to induce different behavioral effects for different illumination conditions (e.g. number of active LEDs per illumination condition, and the minimum distance between LEDs across illumination conditions) is critically dependent on the behavioral task, cortical area, and viral expression levels. Here, we establish the proof of concept for 4 LEDs and 2mm cortical distance. However, future experiments are required to paint the bigger picture.

Together, these results demonstrate the potential utility of Opto-Array for optogenetic perturbation experiments in non-human primates. We note that this tool improves the utility of optogenetics in large brains by advancing on the method of light delivery, and could be further enhanced in the future to include recording probes as well. In sum, Opto-Array offers a chronically implantable solution to the problem of light delivery in optogenetic experiments, particularly for large brains where the problem is pronounced. As such, it may help enable safer, chronically-reproducible behavioral optogenetics experiments in nonhuman primates.

## Methods

### Characterization of Opto-Array

#### Photometric measurements

Photometric measurements were made with a power-meter (Thorlabs) with power sensor in tight proximity (<0.5mm) to the surface of the LED arrays, mimicking the distance between the sutured LED array to the cortical surface. We averaged the power output over a sensor of 9mm in diameter and over a 500ms duration window. To measure the spatial density of LED power, we measured the power output of individual LEDs with the same power-meter, but with an pin-hole occluder placed in between, with varying pin-hole size. In order to mitigate mis-alignments of LEDs with respect to the power sensor, we repeated this experiment with all LEDs on the array and selected the LED with maximally detected power. We additionally repeated this experiment on an Opto-Array that was implanted in an animal for >6months. The light output of the explanted array approximately matched that of a new one (Figure S1D), demonstrating the survivability of this tool in-vivo.

### Temperature measurements

We measured the thermal response of an Opto-Array implanted directly on the cortical surface of an adult rhesus monkey in two separate experiments. Temperature was sampled from the embedded thermistor every 30ms. We note that this measurement is a highly conservative upper bound for the corresponding temperature change on the cortical surface, given the silicone insulation that separates the thermistor from the brain. Under a simplified model of heat transfer (assuming specific heat capacity ranging between 0.2 and 2.55 W/m.K for the silicone, and 0.3 W/m.K for the PCB), we expect an increase of only 0.03° to 0.26° on the external surface of the Opto-Array for every 1° increase measured by the thermal sensor. It is also worth mentioning that temperature readings vary depending on the distance of each LED from the thermistor on the PCB. To factor out the apparent thermal effect of LED distance from the thermistor we used only the LEDs that are adjacent to the thermistor. To ensure the animal’s safety, in both experiments, trials in which the PCB temperature increased more than 3^°^C were aborted.

In experiment 1, we measured the LED thermal response after a single activation. Each trial lasted for 11 seconds and contained one activation that started 1s after the onset of the trial. Each activation condition was randomly selected from a set of combinatory conditions including the following parameters: the number of active LEDs (1, 3 or 5), duration of activation (100, 200 or 500ms), power of activation (0, 40, 82, or 132mW). Each trial-type was repeated 10 times, except for the trials in which the temperature crossed the 3^°^C safety limit (see Figure S1C).

In experiment 2, we measured the thermal response during sequences of LED activations. Each trial started with recording 1 second of baseline temperature prior to sequences of LED activations that lasted each 10 minutes. Each activation sequence was randomly selected from a set of 40 combinatory conditions including the following parameters: the number of active LEDs (1 or 5), duration of activation (200ms or 500ms), power of activation (82 or 191mW) and duty cycle of activation (one pulse every 1, 2, 4, 8 or 16 seconds).

### Behavioral effects of optogenetic perturbation

#### Subjects and surgery

Behavioral data were collected from one adult male rhesus macaque monkey (Macaca mulatta, subject Y). Monkey Y was trained on a two-alternative forced-choice luminance discrimination task (Figure 2A). Following this, we injected AAV8-CAG-ArchT on the right hemisphere of the primary visual (V1) cortex, covering a region of 15mmx7mm with over 18 injection sites, injecting 3ul at a rate of 200nl/min in each each site (described in detail in Open optogenetics). Over this transfected tissue, we first implanted a steel recording chamber (Crist) for acute optrode experiments, and confirmed the viral expression by recording modest neural modulation by delivery of green light (Figure S1A). We did this to confirm viral expression using a traditional method, but typically this stage is not typically needed and we recommend covering the viral injection zone with artificial dura before closing the dura on it. The layer of artificial dura (between pia and dura) prevents tissue adhesions and makes the second surgery smoother. In a second surgery, we removed the chamber and implanted two 5×5 LED arrays over the transfected tissue. To provide access for array implantation, a large U shaped incision (5mmx10mm, base of the U being the long side) was made in dura mater. Viral expression can also be confirmed at this stage using an alternative method: looking for fluorescence produced by GFP. After opening the dura the lights of the operating room can be turned off, then using a flashlight with appropriate wavelength and proper goggles (e.g. 440-460nm excitation light, 500nm longpass filter for GFP) the fluorescence of the viral expression zone can be directly inspected and photographed. Besides confirming the viral expression, one advantage of this method is to visualize the expression zone and implant the array precisely over it. The array was kept in position by suturing the holes in the corners of the arrays to the edges of the rectangular opening in the dura (using non-absorbable suture). This tightly keeps the arrays aligned with the pia surface directly under them. The dura flap was loosely sutured over the arrays (to avoid putting pressure on the cortex) and the area was covered with DuraGen. Schematics of this surgical procedure are shown in Figure 1F. All procedures were performed in compliance with National Institutes of Health guidelines and the standards of the MIT Committee on Animal Care and the American Physiological Society.

### Behavioral paradigm

The luminance discrimination behavioral task was designed to probe the role of millimeter scale regions of V1, which encode local features of the visual field. Stimuli were presented on a 24’’ LCD monitor (1920 × 1080 at 60 Hz; Acer GD235HZ) and eye position was monitored by tracking the position of the pupil using a camera-based system (SR Research Eyelink 1000). At the start of each training session, the subject performed an eye-tracking calibration task by saccading to a range of spatial targets and maintaining fixation for 800 ms. Calibration was repeated if drift was noticed over the course of the session.

Figure 2A illustrates the behavioral paradigm. Each trial of the behavioral task consisted of a central visual fixation period, during which the animal had to hold gaze fixation on a central fixation spot for 900ms. During this epoch, one (or none) of the LEDs were pre-emptively activated on a random proportion of trials. This was followed by the simultaneous and brief (50ms) presentation of two sample stimuli (Gaussian blob of 1 degree size, varying in luminance) in the periphery, at radially opposite locations in the visual field. The LED activation was timed to completely overlap the stimulus-related activity in V1. Following the extinction of these stimuli, two target dots were presented at the stimulus locations. The task required the subject to make a saccade to a target location defined by the brighter of the two sample stimuli. By varying the relative luminance of the two sample stimuli, we systematically varied the task difficulty. Correct reports were rewarded with a juice reward. Real-time experiments for monkey psychophysics were controlled by open-source software (MWorks Project http://mworks-project.org/).

### Optical fiber experiments

To provide a baseline for comparison across methodologies, we first performed a small number of acute optical fiber experiments. We first confirmed weak viral expression by recording modest neural modulation by delivery of green light via an acutely inserted optical fiber (Figure S1A). Next, we measured the behavioral effects of optogenetic suppression with light delivered via an acutely inserted fiber. Figure S1B shows the behavioral effects in the two alternative forced choice luminance discrimination task described above, for an example optrode session. Formatting is as in Figure 2B.

### Opto-Array experiments

Behavioral data with LED activation was collected over NN behavioral sessions, with NN+NN (mean + SD) trials per session. For the first set of experiments, we activated groups of four neighbouring LEDs simultaneously to increase both the spatial spread and power of light. We interleaved four such groups, each consisting of four corners of arrays. Given the chronic nature of this tool, we collected behavioral data over several sessions while activating LEDs on a small (20%) portion of trials, with the same illumination (900ms) duration that yielded neural suppression and behavioral effects in optrode experiments.

### Behavioral analysis

To assess the behavioral effects from stimulation, we fit psychometric functions to the animal’s behavioral choices, separately for each LED condition (including the control condition of no LED illumination), and for each tested position in the visual field. For each tested location (parameterized in polar coordinates with r, θ), we pooled all trials where either of the target or distractor stimuli were presented in a pooling region spanning 4° along the radial dimension and *π/8* along the angular dimension. For this subset of trials, we fitted a psychometric curve for each LED condition using logistic regression:

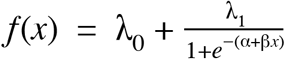

Where λ_0_, λ_1_, α, β are the fitted parameters and *f* (*x*), *x* correspond to the dependent and experimentally controlled variables. *x* corresponds to the visual signal, the difference in luminance between the stimulus in the pooling region and the stimulus outside the pooling region, on each trial. *f* (*x*) models the choice, 1 for choice in the pooling region, 0 for choice outside the pooling region, on each trial. λ_0_, λ_1_ model lapses, i.e. the floor and ceiling values of the psychometric function, attributed to visual deficits not resulting from LED illumination. α, β model the criterion and sensitivity of the psychometric function. We fit psychometric functions with constrained non-linear least squares using standard Python libraries (scipy.curve_fit) and extracted both the fitted parameter estimates (e.g.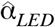) and the variance of parameter estimates (e.g 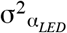). Note that psychometric functions were fit to individual trial data, such that the variance in the parameter estimates captures trial-by-trial variability.

To assess the effect of LED activation, we measured the change in psychometric criterion (i.e. corresponding to shifts in the psychometric curves) via the difference in estimated criterion between the function fits of the LED condition and the control condition: 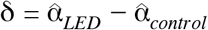. We normalized this difference by the pooled variance 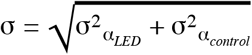 to obtain a z-scored metric: 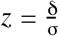. Repeating this procedure for each tested location in the visual field, we obtained a 2D map of z-scored psychometric shift estimates. Z-scores were converted to one-tailed p-values using the survival function of the normal distribution *N(0,1)*.

We used a region of interest (ROI) based on the functional organization of primate V1: the dorsal region of V1 on the right hemisphere is known to represent the contralateral (left) lower visual field. Given that viral expression in monkey Y was verified to be poor and likely inhomogeneous over the cortical tissue, we did not attempt to localize behavioral effects from LED illumination with finer precision.

## Author Contributions

M.S. designed and fabricated the Opto-Array, with guidance from A.A. and J.J.D.

R.R. performed the Opto-Array photometric experiments.

S.B. and R.A. performed the Opto-Array thermal experiments, with guidance from A.A..

R.R. and A.A. performed the optical fiber experiments, with guidance from J.J.D.

R.R. performed the Opto-Array behavioral experiments, with guidance from A.A. and J.J.D.

R.R. and A.A. wrote the manuscript. All authors reviewed the manuscript.

## Acknowledgements

We thank Edward S Boyden and Jacob G Bernstein for their critical help during the early stages of array development. This research was supported by Simons Foundation (SCGB [325500] to JJD), US National Eye Institute grants R01-EY014970 (to J.J.D.), and NIH Grants K99 EY022924 (to A.A.) and by the Intramural Research Program of the NIMH ZIAMH002958 (to A.A.).

## Supplemental Information

**Figure S1.**
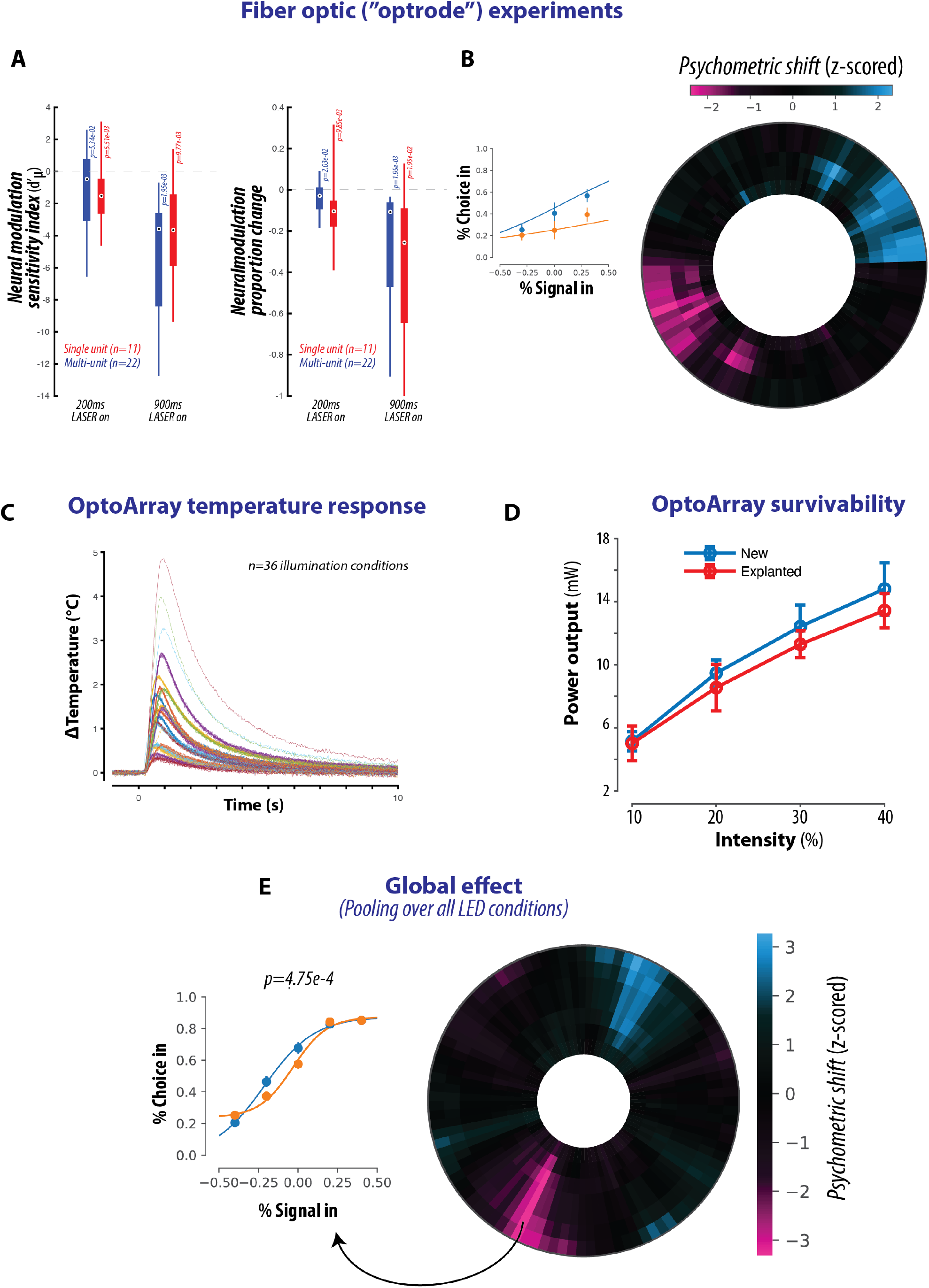
(a,b) Results from fiber optic experiments. (a) After injection of AAV8-CAG-ArchT on the right hemisphere of the primary visual (V1) cortex, we first implanted a steel recording chamber (Crist) for acute optrode experiments. We recorded V1 responses to a brief full-field grating stimulus, interleaving trials with and without light delivery from the acutely inserted optic fiber coupled to a green light LASER. We confirmed weak viral expression: over all recorded neural sites, the neural modulation (silencing) by light delivery was poor but significant, as quantified by the sensitivity (d’ between control and light trials) and the proportion of silenced evoked spikes. (b) Next, we measured the behavioral effects of optogenetic suppression with light delivered via the acutely inserted fiber. The behavioral effects in the two for an example fiber optic session is shown, with formatting is as in Figure 2B. We observe significant psychometric shifts in the region of interest within the visual field. (c) Average thermal response from implanted Opto-Array to 36 different LED conditions, varying in power, duration, and number of illuminated LEDs. (d) Survivability test comparing the light power output of new Opto-Array to one explanted from an animal. (e) Global effect from Opto-Array experiments. Pooling over all LED conditions and over the entire ROI, we observed a reliable behavioral effect of LED illumination even at this coarse scale, in the form of a statistically significant psychometric shift away from the ROI (p=4.75e-4, Figure S1E).

